# Plasma proteome profiling for AKI biomarker candidates associated with first-line ART in people living with HIV in South Africa

**DOI:** 10.1101/2024.07.26.605118

**Authors:** Rethabile J. Mokoena, Sylvia Fanucchi, Firdaus Nabeemeeah, Ebrahim Variava, Neil Martinson, Ireshyn S. Govender, Previn Naicker

## Abstract

With the highest global burden of HIV, South Africa initiates HIV treatment using first-line antiretroviral therapy (ART) regimens in newly diagnosed patients. Antiretroviral (ARV)-associated nephrotoxicity has been observed in ~10% of South African patients newly receiving first-line ART, as well as in patients who have extensively received TDF-based ART regimens, and this can progress to acute kidney injury (AKI). To identify potential biomarkers that will improve the detection of ARV-associated nephrotoxicity, proteomic analysis was performed using the sequential window acquisition of all theoretical mass spectra (SWATH-MS) data acquisition method on the plasma of the case group (AKI) and the control group (non-AKI). Data are available via ProteomeXchange with identifier PXD054218. Evaluation of the results identified thirty-four proteins that showed a significant change in abundance, with three proteins showing increased abundance, while thirty-one proteins showed decreased abundance between the AKI and non-AKI groups. Machine learning was also used to evaluate the results and showed twenty ranked proteins that contributed to distinguishing between the AKI and non-AKI groups. The majority of the proteins of significant differential abundance and those ranked by the machine learning model participate in enriched biological processes that correspond to known pathophysiological and cellular mechanisms that contribute to AKI, suggesting that those proteins can serve as potential biomarkers to be further verified and validated for AKI diagnosis. Comparison of the plasma and previously generated urinary proteome profiles showed five proteins with significant differences in abundance that overlapped both proteomes providing evidence for the renal physiological dynamics of these proteins in renal injury and disease.

*Significance*: Improving the detection of ARV-associated nephrotoxicity is challenging because current serum creatinine (srCr) based-equations used to estimate the glomerular filtration rate (eGFR) for assessing kidney function have been reported to overestimate the eGFR in the South African population and changes in srCr measurements also reflect a delayed response to the initial structural injury of the kidney. This hinders the timeous detection of ARV-associated nephrotoxicity which can lead to irreversible renal damage and can compromise the continuation of treatment in PLHIV, thus emphasising the need to introduce novel AKI biomarkers to mitigate ARV-associated nephrotoxicity in at risk PLHIV in South Africa.

**Graphical Abstract:** 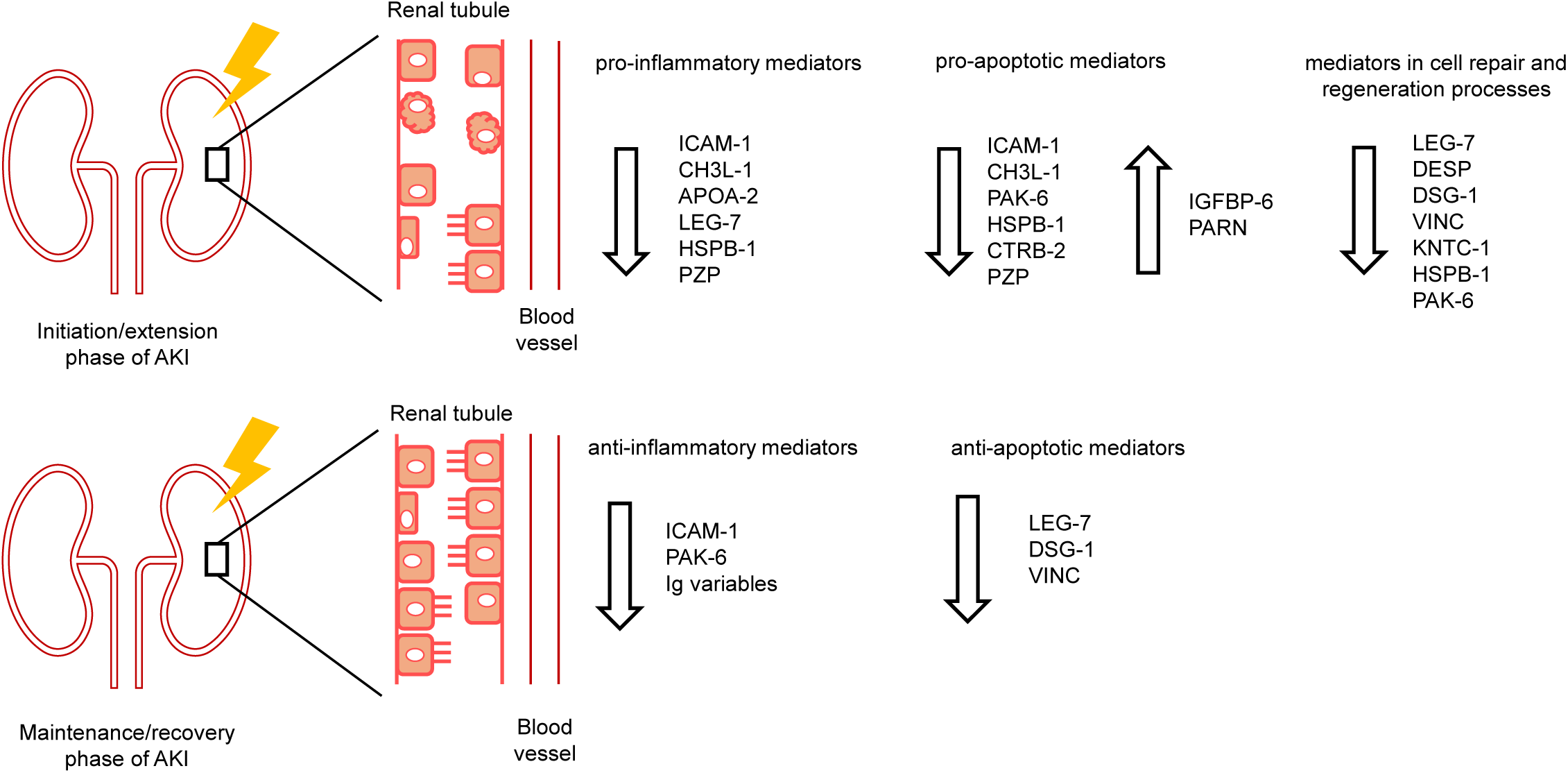

**Highlights:**

- Identified thirty-four proteins with significant changes in abundance between study groups.
- Machine learning showed fourteen proteins associated with renal disease biology that differentiate the study groups.
- First study to analyse plasma proteome of this cohort of PLHIV in South Africa
- Five proteins overlap the plasma and urinary proteome profiles of the study groups.

## 1. Introduction

South Africa (SA) is the epicentre of the HIV/AIDS pandemic, with ~19.6% of the adult population living with HIV (PLHIV) [1]. SA also hosts the largest antiretroviral therapy (ART) program worldwide, initiating treatment with the first-line ART regimens consisting of tenofovir disoproxil fumarate (TDF) and providing free access to ART for - 5.5 million PLHIV [1, 2, 3]. Approximately 10% of South African patients newly receiving first-line ART experience acute adverse nephrotoxicity [4] and extensive clinical administration of TDF-based regimens can cause nephrotoxicity [5, 6]. ARV-associated nephrotoxicity can progress to acute kidney injury (AKI) [5, 6], which is characterised by the dysfunction of the proximal tubule of the kidney [7, 8, 9]. Improved detection of ARV-associated nephrotoxicity encourages ART adherence and continuation by substituting the afflicting ARV drug, ensuring that irreversible renal damage is mitigated [5, 10].

However, early detection of ARV-associated nephrotoxicity is challenging because patients can be asymptomatic or experience mild to moderate nephrotoxic events that can go undetected due to the invalidity of current glomerular filtration rate (GFR)-estimating equations that overestimate GFR in African patients [6, 11]. Additionally, serum creatinine (srCr) measurements are unreliable for early detection because they reflect a delayed response to the structural changes occurring at the onset of AKI [12, 13, 14, 15]. This results in misinformation regarding the state of kidney function in African patients, allowing for unimpeded progression to irreversible renal damage [5, 6, 10, 11].

Mass spectrometry (MS)-based proteomics can be used to detect changes in protein abundances associated with ARV-associated AKI [16, 17, 18]. These changes are often related to the onset and progression of a disease, enabling the identification of diagnostic biomarkers that can provide insight into the underlying pathophysiological mechanisms of ARV-associated AKI [18, 19, 20, 21, 22]. Although novel biomarkers for AKI have been identified, few have been introduced for the diagnosis of AKI [13, 14, 23]. This emphasises the need for new AKI biomarkers, especially within the South African context, that are sensitive and specific for identifying the onset and progression of nephrotoxicity in PLHIV. This study aimed to identify and quantify differentially abundant plasma proteins that can distinguish between South African PLHIV clinically diagnosed with AKI versus those with normal kidney function to aid in the improved detection of ARV-related nephrotoxicity.

## 2. Material and Methods

### 2.1) Ethics Statement

Ethics clearance for this research project was obtained from the Research Ethics Committee of the Council of Scientific and Industrial Research in Pretoria, South Africa (#58/2013) and the Human Research Ethics Committee of the University of the Witwatersrand in Johannesburg, South Africa (#120612). Patient recruitment and sample collection was conducted at the Klerksdorp Tshepong Hospital, in the North West Province of South Africa with informed written consent from enrolled patients.

### 2.2) Sample collection and study cohort

Blood samples were collected by venipuncture from HIV positive patients on TDF-ART regimens in the adult (≥18 years) wards by trained clinical staff into EDTA collection tubes. Individual blood samples were centrifuged at 1 000 x g for 10 minutes at 4°C to separate the cellular components of blood from plasma. The plasma tubes were centrifuged at 2 500 x g for 10 minutes at 4°C to ensure that all cellular blood contaminants were removed from the plasma and crude plasma samples were stored in vials as 1 ml aliquots at -80°C for analysis. One hundred and thirty-five patient plasma samples were analysed in this AKI study, with 53 patients in the AKI (case) group and 82 patients in the non-AKI (control) group based on their kidney function according to the 2014 Clinical Practice Kidney Disease Improving Guidelines Outcome (KDIGO), where AKI was diagnosed in patients showing increased srCr ≥ 0.3 mg/dL within 48 hrs or 1.5-times the srCr baseline that was determined or estimated in the last 7 days. Patients belonging to the case group were matched according to age (± 3 years), gender, and race, to patients in the control group. The exclusion criteria were patients with pre-existing renal complications or prior renal injury, as well as patients who were unwilling to have follow-up visits and patients that had medical records lacking sufficient information for determining any prior use of HIV treatment.

### 2.3) Sample preparation

To prepare crude plasma samples for analysis, 20 μl of plasma was diluted (25x) with buffer (50 mM Tris-HCl, 2% SDS; pH 8) and 20 μg of each diluted sample (as per Bicinchoninic Acid (BCA) Protein Assay, Sigma-Aldrich, Missouri, USA) was reduced using dithiothreitol (DTT) (10 mM *v*/*v*, 30 min, 60°C) and alkylated with iodoacetamide (IAA) (40 mM *v*/*v*, 30 min, RT-dark) (Sigma-Aldrich, Missouri, USA), and quenched with DTT (10 mM *v*/*v*). Protein digestion and hydrophilic interaction liquid chromatography (HILIC) sample clean-up was performed as per the HILIC protocol described in Baichan *et al* (2023) [81], using the KingFisher Flex^TM^ (Thermo Scientific^TM^, Massachusetts, USA), an automated magnetic handling station. Briefly, the MagReSyn® HILIC microspheres (ReSyn Biosciences, Edenvale, South Africa) were equilibrated in equilibration buffer (100 mM ammonium acetate (NH_4_Ac), 15% acetonitrile (ACN); pH 4.5), followed by protein binding onto the microspheres with a protein:microsphere ratio of 1:10. Thereafter, two washes of 95% ACN were performed, followed by on-bead protein digestion with trypsin (Promega, Wisconsin, USA) and endopeptidase LysC (Thermo Scientific^TM^, Massachusetts, USA) (1:20 and 1:100 protease:protein, respectively) in 50 mM ammonium bicarbonate for 2 hours, at 47°C. Digestion was stopped using 1% trifluoroacetic acid (TFA) and the resulting peptides were collected using a magnetic rack and transferred into protein Lo-Bind Eppendorf tubes (Eppendorf, Hamburg, Germany). The peptides were stored at - 80°C and were dried using a vacuum concentrator (CentriVap® Concentrator, Labconco, Missouri, USA), and were resuspended in 2% ACN, 0.2% formic acid (FA), *v*/*v* for LC-MS/MS analysis.

### 2.4) Liquid chromatography-tandem mass spectrometry (LC-MS/MS) data acquisition

Resuspended peptides (0.5 μg per sample as per the Pierce Quantitative Colorimetric Peptide Assay, Thermo Scientific^TM^, Massachusetts, USA) were loaded into Evotips as per manufacturer’s instructions and were injected into the Evosep One LC system (EV-1000, Evosep Biosystems, Odense, Denmark). Peptides were then separated on a C-18 Evosep performance column (EV-1109: 8 cm, 150 pm x 1.5 pm) at a flow rate of 1 pl/min. The generic 60 SPD method had peptides eluted over a 21-minute gradient consisting of a preformed gradient of 5 - 30% Solvent B and an offset gradient with a lower percentage of ACN using Solvent A (Solvent A: LC-MS grade water with 0.1% FA; Solvent B: 100% LC-MS grade ACN with 0.1% FA) (Thermo Scientific^TM^, Massachusetts, USA). The Evosep One LC system was coupled to a TripleTOF 5600 mass spectrometer (SCIEX, Framingham, Massachusetts, USA) set in a positive mode. The instrument was operated using the sequential window acquisition of all theoretical fragment ion spectra (SWATH) acquisition method, consisting of 48 variable windows (VW) with 1 m/z overlap between VW, and measured 48 MS/MS scans as well as a single MS scan over a mass range of 400 - 1 100 m/z. The accumulation time for the MS and MS/MS scans was 50 and 20 milliseconds (msec), respectively, with a total cycle time of 1 062 msec.

### 2.5) LC-MS/MS data processing using Spectronaut^TM^ 17

The SWATH-MS raw data were analysed with Spectronaut^TM^ 17 (Biognosys AG, Schlieren, Switzerland), using the directDIA+ workflow with the default settings applied: specific enzyme cleavage by trypsin/P and LysC/P allowing for 2 consecutive missed cleavage sites, fixed modification was set for cysteine carbamidomethyl, while variable modifications were set for acetyl at protein N-terminal and methionine oxidation, identifications were performed at a false discovery rate (FDR) of 1% at a peptide and protein group level. For quantification, no imputation strategy was applied, the protein LFQ method applied was automatic, quantity type was set to area under the curve (AUC), and quantification was based on MS/MS (fragment ion) measurements. A search database consisting of all reviewed human protein FASTA sequences including contaminant proteins from Uniprot Knowledge Database was used for peptide identification and protein inference. Protein inference was performed by the IDPicker inference algorithm [65], and the study groups (AKI or non-AKI) were specified.

### 2.6) Data analysis and machine learning

Unpaired t-tests in Spectronaut^TM^ 17 were performed to compare protein group quantities in the AKI and non-AKI groups. The average protein intensities were log2 transformed and identification was performed at a protein q-value cutoff of 5% for each run and 1% across the experiment. The p-values were corrected for the multiple hypothesis testing by controlling the false discovery rate (FDR) [66, 67]. A retrospective power analysis was performed to determine a desired fold-change at a power of 80%, with an FDR of 1%, and a conservative sample size of 53 patient samples using MSstats (v3.4.1) [25] in RStudio (v2022.12.0 + 353) (R Core Team, 2022). The selection criteria used to identify significantly differentially abundant proteins included q-value < 0.05 and fold-change > 1.68 as per the retrospective power analysis.

OmicLearn (v1.4) was used to perform analysis of proteomic data by exploring machine learning (ML) models. Within OmicLearn the feature tables were imported using the Pandas package (v1.0.1) and manipulated using the NumPy package (v1.18.1). The ML pipelines were built using the scikit-learn package (v0.22.1) and ML was completed in Python (v3.8.8) [26, 27].

### 2.7) Bioinformatics and statistical analysis

Patient demographic and clinical characteristic data were analysed using non-parametric statistical tests in Epitools, an online platform for epidemiological and clinical statistics [28], with a p-value cut-off < 0.05. The Chi-squared test of independence was used to compare sex and race between the AKI and non-AKI groups, respectively. The Mann-Whitney U Test (Wilcoxon signed rank test) was performed to compare the median age, srCr levels, and the eGFR between the AKI and non-AKI groups, respectively.

Protein abundance data was analysed using Enrichr [29, 30] for functional enrichment analysis. The enriched pathways and gene ontology processes and functions were ranked by p-value and adjusted p-value from most to least significant, with a p-value cut-off of 0.05.

Significantly enriched GO processes, functions, and pathways were reported. The volcano plot was generated using SRplot, an online platform for data analysis and visualisation [31].

## 3. Results

### 3.1) Overview of AKI cohort

Briefly, 135 plasma samples from HIV-positive patients on TDF-ART regimens were collected at a single-time point, with 53 and 82 plasma samples in the AKI and non-AKI study groups, respectively. Statistical analysis of the race, sex and the median age of the patients showed no significant difference (p > 0.05), whereas the median eGFR and srCr levels of the patients showed a significant statistical difference (p ≤ 0.05) between the AKI and non-AKI groups, with an elevation in the srCr levels and a decline in the eGFR in the AKI group (Table 1).

**Table 1:** Demographic details and AKI clinical characteristics of study participants in the AKI and non-AKI groups. The demographic details consist of the sex, median age, and race/ethnicity of study participants. The median clinical characteristics (eGFR and srCr levels) of the patients are also represented in the table.

| Category | Normal ranges | AKI (cases)<br>n = 54<br>Median<br>[IQR] | non-AKI (controls)<br>n = 82<br>Median<br>[IQR] | p-value |
| --- | --- | --- | --- | --- |
| **Sex:** |  |  |  | **0,45** |
| **Females, n (%)** | - | 32 (59,26%) | 55 (67,07%) |  |
| **Males, n (%)** | - | 22 (40,74%) | 27 (32,93%) |  |
| **Age:** |  |  |  | **0,37** |
| **In years** | - | 42<br>[36,25 - 51,75] | 40,5<br>[35 - 52,75] |  |
| **Race:** |  |  |  | **0,44** |
| **Black, n (%)** | - | 52 (96,30%) | 81 (98,78%) |  |
| **Coloured, n (%)** | - | 1 (1,85%) | 1 (1,22%) |  |
| **Indian, n (%)** | - | 1 (1,85%) | 0 |  |
| **Serum creatinine levels (µmol/L):** |  |  |  | **< 0,0001** |
|  | 53 - 97,2<br>(females) | 276<br>[146,25 - 520] | 67<br>[58 - 81,5] |  |
|  | 61,9 - 114,9<br>(males) |  |  |  |
| **Estimated glomerular filtration rate (mL/min/1.73m<sup>2</sup>):** |  |  |  | **< 0,0001** |
|  | ≥ 90 | 21,25<br>[9,93 - 35,21] | 82,03<br>[68,58 - 92,88] |  |
| The Chi-square Test of Independence was performed on the sex, and race. The Mann-Whitney U Test (Wilcoxon signed rank test) was performed on data for the median age, eGFR and srCr levels.<br>Data for normal ranges indicated were retrieved for the serum creatinine levels [32] and for the estimated glomerular filtration rate levels [33]. |  |  |  |  |

The variation in the LC-MS/MS performance was evaluated to assess the quality and reliability of the experimental data obtained over the course of five days of analysis. Low inter-day variation between the protein groups (Figure S1 and S2) and the peptides (Figure S1) identified in the quality control replicates was observed. Additionally, the peak capacity, median full width half maximum (FWHM) and the average data points per peak indicated that there was adequate peptide peak separation in this study (Table S1). The sequential window acquisition of all theoretical fragment-ion spectra (SWATH) acquisition method employed for proteome profiling resulted in an average of 3 212 modified peptides and 363 protein groups, with an average of 3 365 modified peptides and 380 protein groups identified in the AKI group, while an average of 3 113 modified peptides and 352 protein groups were identified in the non-AKI group (Figure S3).

### 3.2) Plasma proteome differences and machine leaning-based classification between AKI and non-AKI groups

Plasma proteins from the AKI and non-AKI groups were compared using the Student’s t-test, and 34 proteins with significantly altered abundances (i.e. candidate markers) were identified, of which 3 showed increased abundance (pink) and 31 showed decreased abundance (blue) in the AKI group (Figure 1A) using the criterion: q-value ≤ 0.05, log_2_FC > 0.75 (FC > 1.68), as per the retrospective power analysis (Figure S4). Functional enrichment analysis of the candidate markers showed enriched GO biological processes and pathways of signal transduction (e.g., intrinsic/extrinsic apoptotic signalling), immune responses (e.g., leukocyte cytotoxicity), metabolism (e.g., RNA and cholesterol) and small molecule transport (plasma lipoprotein assembly/remodelling) (Table 2).

**Figure 1:**
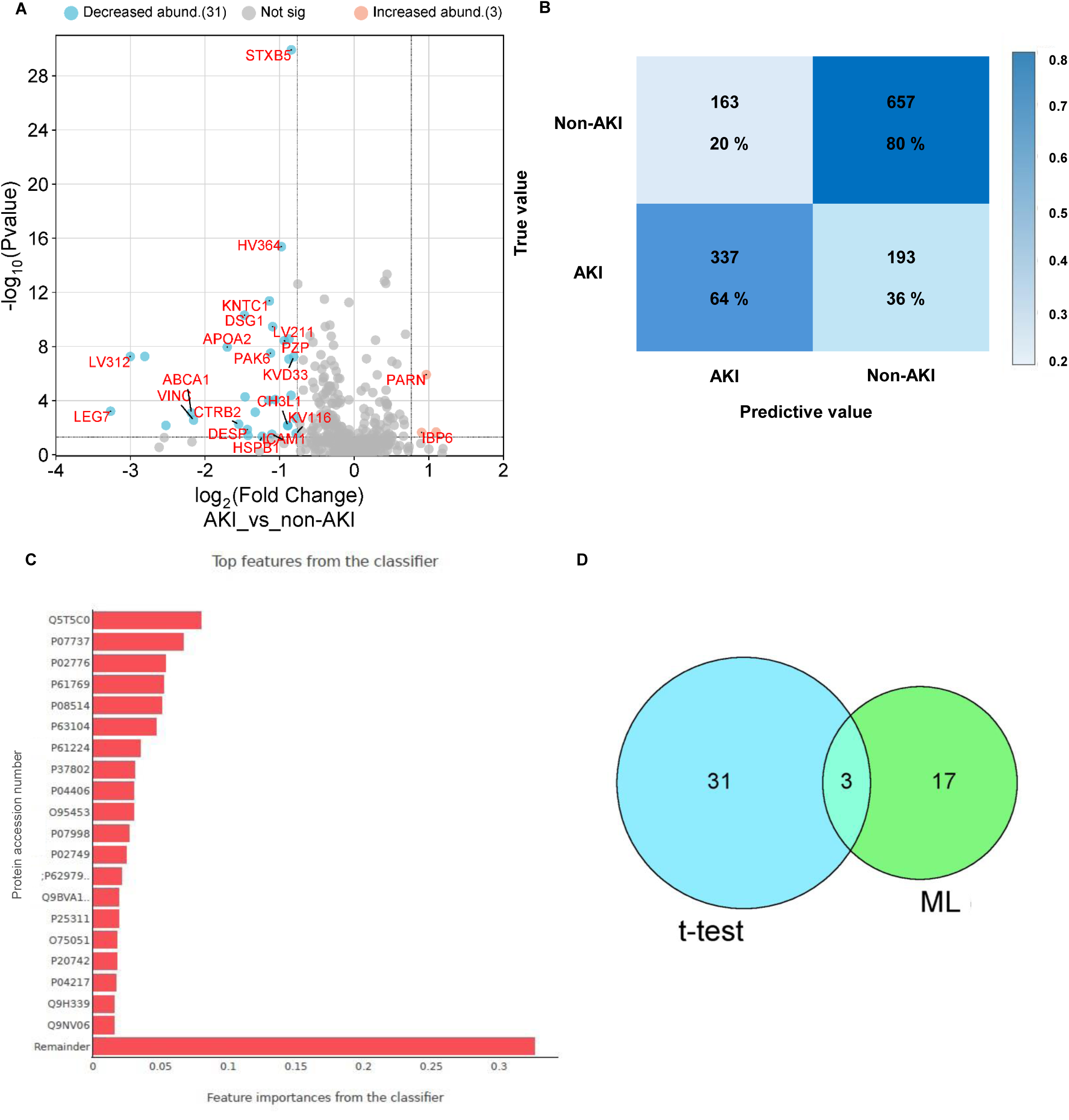
Results of plasma proteome differences and machine learning classification between the AKI and non-AKI groups. A) Volcano plot of differentially abundant proteins generated from the AKI study. B) Confusion matrix of the best performing machine learning (ML) model. C) List of top 20 proteins differentiating the two study conditions ranked in order of importance using machine learning (ML). D) Venn diagram comparing differentially abundant proteins identified by an unpaired t-test and important proteins used by the machine learning (ML) model in B).

**Table 2:** GO biological processes, molecular functions, and pathway analysis of candidate markers. Enriched GO biological processes, molecular functions, cellular compartments as well as KEGG, Reactome, and Wikipathways associated with the identified candidate markers is tabulated below. Data generated with Enrichr and is tabulated in descending order of the number of candidate markers found in each process, function, or pathway name (adjusted p-value < 0.05).

|  | Name | Candidate marker gene names | p-value | Adjusted p-value |
| --- | --- | --- | --- | --- |
| **GO Biological Processes** | Regulation Of Very-Low-Density Lipoprotein Particle Remodelling (GO:0010901); Regulation of CoA-transferase Activity (GO:1905918); Regulation of Cholesterol Transport (GO:0032374); Regulation of Lipid Catabolic Process (GO:0050994); Regulation of Cytokine Production Involved in Immune Response (GO:0002718) | *APOA2* | 8,72 E-03 | 1,32 E-01 |
|  | Positive Regulation of Cell Migration by Vascular Endothelial Growth Factor Signalling Pathway (GO:0038089); Vascular Endothelial Growth Factor Receptor Signalling Pathway (GO:0048010) Regulation of Cytokine Production Involved in Immune Response (GO:0002718) | *HSPB1* | 1,22 E-02 | 1,32 E-01 |
|  | Regulation Of Oxidative Stress-Induced Intrinsic Apoptotic Signalling Pathway (GO:1902175); Negative Regulation of Oxidative Stress-Induced Cell Death (GO:1903202) | *HSPB1* | 3,10 E-02 | 1,32 E-01 |
|  | Regulation Of Leukocyte Mediated Cytotoxicity (GO:0001910); Negative Regulation of Extrinsic Apoptotic Signalling Pathway Via Death Domain Receptors (GO:1902042); Negative Regulation of Endothelial Cell Apoptotic Process (GO:2000352); T Cell Activation Involved in Immune Response (GO:0002286) | *ICAM1* | 1,56 E-02 | 1,32 E-01 |
|  | Spindle Assembly Checkpoint Signalling (GO:0071173) | *KNTC1* | 4,45 E-02 | 1,32 E-01 |
|  | RNA Phosphodiester Bond Hydrolysis (GO:0090501) | *PARN* | 4,79 E-02 | 1,32 E-01 |
| **GO Molecular Function** | poly(A)-specific Ribonuclease Activity (GO:0004535); Telomerase RNA Binding (GO:0070034) | *PARN* | 2,08 E-02 | 2,06 E-01 |
|  | Peptidase Inhibitor Activity (GO:0030414); Endopeptidase Regulator Activity (GO:0061135); Endopeptidase Inhibitor Activity (GO:0004866) | *PZP* | 6,28 E-02 | 2,09 E-01 |
|  | Serine-Type Endopeptidase Activity (GO:0004252); Serine-Type Peptidase Activity (GO:0008236); Endopeptidase Activity (GO:0004175) | *CTRB2* | 1,97 E-01 | 3,59 E-01 |
| **GO Cellular Compartment** | Desmosome (GO:0030057) | *DSG1* | 2,93 E-02 | 1,42 E-01 |
| **KEGG Pathways** | Cholesterol metabolism (map04979) | *ABCA1; APOA2* | 3,46 E-03 | 1,21 E-01 |
|  | T cell receptor signalling pathway (map04660) | *PAK6* | 1,67 E-01 | 2,89 E-01 |
|  | NF-kappa B signalling pathway (map04064); TNF signalling pathway (map04668); Natural killer cell mediated cytotoxicity (map04650) | *ICAM1* | 1,67 E-01 | 2,89 E-01 |
| **Reactome Pathways** | Regulation Of mRNA Stability by Proteins That Bind AU-rich Elements R-HSA-450531 | *PARN; HSPB1* | 1,04 E-02 | 3,43 E-01 |
|  | HDL Assembly R-HSA-8963896; Plasma Lipoprotein Assembly R-HSA-8963898 | *ABCA1* | 1,22 E-02 | 3,43 E-01 |
|  | Uptake Of Dietary Cobalamins into Enterocytes R-HSA-9758881; Cobalamin (Cbl, Vitamin B12) Transport and Metabolism R-HSA-196741; Activation of Matrix Metalloproteinases R-HSA-1592389 | *CTRB2* | 1,74 E-02 | 3,43 E-01 |
|  | Apoptotic Cleavage of Cell Adhesion Proteins R-HSA-351906; Apoptosis R-HSA-109581; | *DSG1* | 1,91 E-02 | 3,43 E-01 |
|  | KSRP (KHSRP) Binds and Destabilizes mRNA R-HSA-450604; Deadenylation Of mRNA R-HSA-429947; ATF4 Activates Genes in Response to Endoplasmic Reticulum Stress R-HSA-380994 | *PARN* | 2,93 E-02 | 3,43 E-01 |
|  | Interleukin-10 Signalling R-HSA-6783783; Interleukin-4 And Interleukin-13 Signalling R-HSA-6785807; Interferon Gamma Signalling R-HSA-877300 | *ICAM1; CH3L1* | 7,59 E-02 | 3,43 E-01 |
|  | MAPK6/MAPK4 Signalling R-HSA-5687128; VEGFA-VEGFR2 Pathway R-HSA-4420097; Signalling by VEGF R-HSA-194138 | *HSPB1* | 1,46 E-01 | 3,65 E-01 |
| **Wikipathways** | Metabolic Pathway of LDL HDL And TG Including Diseases WP4522; Statin Inhibition of Cholesterol Production WP430 | *ABCA1; APOA2* | 3,98 E-04 | 2,63 E-02 |
|  | Vitamin B12 Metabolism WP1533; Folate Metabolism WP176; Urotensin II Mediated Signalling Pathway WP5158 | *ABCA1; ICAM1* | 7,04 E-03 | 7,75 E-02 |

Additionally, machine learning (ML) was used to identify proteins that can distinguish the AKI group from the non-AKI group. Various ML pipelines were built, and their performance was evaluated to explore the best performing ML model (Figure S5). The best performing ML model consisted of the ‘no imputation’ method, with the ‘ExtraTrees’ feature selection method and XGBoost classifier. This ML model correctly predicted 64% of AKI patients to the AKI group, while 36% of AKI patients were incorrectly assigned to the non-AKI group. Additionally, this model correctly assigned 80% of the non-AKI patients to the non-AKI group, while 20% of the non-AKI patients were incorrectly assigned to the AKI group (Figure 1B). A ranked list of 20 proteins that contributed the most to differentiating the AKI group from the non-AKI group was generated (Figure 1C).

Furthermore, syntaxin-binding protein 5, pregnancy zone protein and poly(A)-specific ribonuclease were overlapping the candidate markers identified using the Student’s t-test and the ranked list of proteins used by the ML model applied to distinguish the AKI group from the non-AKI group (Figure 1D), while 17 of the ranked proteins were not identified as candidate markers by the Student’s t-test.

### 3.3) Comparison of plasma and urinary proteomes of the AKI and non-AKI groups

Previously acquired urinary proteome profiles [24] of the AKI and non-AKI groups were compared with the plasma proteome profiles of the same study groups. Galectin 7 (LEG-7), chitinase 3-like protein 1 (CH3L-1), insulin-like growth factor-binding protein 6 (IGFBP-6), heat shock protein 0 1 (HSPB-1), and immunoglobulin kappa variable 1D-33 (KVD-33) overlapped the urinary and plasma proteomes of the study groups (q-value < 0.05, log_2_FC > 0.58; FC > 1.5).

The fold-change of LEG-7 was 3.55-times less in plasma than in urine and was greater in the non-AKI group than the AKI group. The fold-change of CH3L-1 was 1.99-times greater in urine than in plasma and the fold-change of CH3L-1 in urine was greater in the AKI group, but in plasma was greater in the non-AKI group. The fold-change of IGFBP-6 was 1.29-times greater in urine than in plasma and was greater in the AKI group than in the non-AKI group. The fold-change of HSPB-1 was 0.83 times greater in plasma than in urine and was greater in the non-AKI group than the AKI group. The fold-change of KVD-33 was 1.46-times greater in urine than in plasma and was greater in the non-AKI group than the AKI group (Table 3).

**Table 3:**
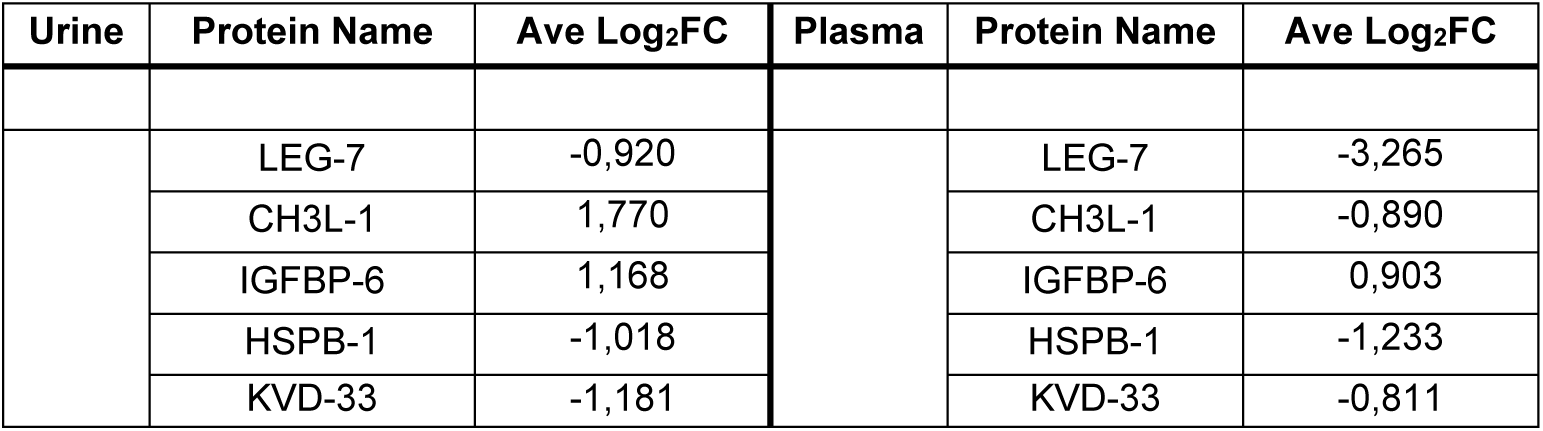
List of candidate markers overlapping in the urinary and plasma proteomes of the study groups. The average log2 fold-change of LEG-7, CH3L-1, IGFBP-6, HSPB-1, and KVD-33 in urine and in plasma.

## 4. Discussion

Plasma proteome changes were evaluated in PLHIV that were clinically diagnosed with AKI and those that were not, to identify differentially abundant proteins between the AKI and non­AKI group. No significant statistical difference was observed in the confounding demographic factors associated with this cohort, while a statistically significant difference was observed in the median srCr levels and eGFR, confirming that patients with kidney dysfunction (AKI group) had elevated srCr levels and lower eGFR compared to patients without kidney dysfunction (non-AKI group), as supported by the literature [12, 13, 14, 15] (Table 1).

Thirty-four differentially abundant protein groups were identified as significant, i.e., candidate markers between the AKI and non-AKI groups. The candidate markers identified (Figure 1 A) were reported to be associated with renal damage and repair processes; however, to date, this study has not been undertaken within this South African population. These dysregulated candidate markers were associated with four main biological processes that are key mechanisms of AKI pathophysiology: immune response, programmed cell death, plasma lipoprotein assembly and cholesterol/lipid metabolism, and processes of cell survival, repair, and regeneration. AKI is usually marked by these four phases: the initiation phase reflects injury of the renal tubular epithelial cells (RTECs), the extension phase reflects persistent injury and inflammatory responses that induce cell death, the maintenance phase initiates cellular repair in damaged but viable cells, and the recovery phase reflects the continuation of restoring normal cellular and organ functions [34, 35, 36], and these four main biological processes play a role in these phases.

### Immune response

The biological processes and pathways of the innate and adaptive immune responses were shown to be enriched in this study, inducing pro- and anti-inflammatory pathways [37, 38]. PAK-6 and ICAM-1 participate in enriched T-cell receptor signalling and activation pathways, as well as the regulation of leukocyte-mediated cytotoxicity (Table 2), and have decreased abundance in the AKI group, suggesting the recovery from AKI [37, 38]. ICAM-1 and CH3L-1 participate in enriched anti-inflammatory pathways involving IL-4, IL-10, and IL-13 signalling (Table 2), and their decreased abundance in the AKI group suggests a progression of AKI [37, 38]. AKI is also associated with the increased production and reduced clearance of cytokines [37, 38]. APOA-2, LEG-7, CH3L-1, and HSPB-1 are involved in enriched pathways that regulate cytokine production, ICAM-1 was shown to participate in enriched TNF signalling pathways, and PZP was associated with clearance of pro-inflammatory cytokines (Table 2), these candidate markers have decreased abundance within the AKI group reflecting a progression of AKI. Furthermore, the Ig-lambda, -kappa, and -heavy variables (Table S5) responsible for antigen recognition in adaptive immune responses, have decreased abundance in the AKI group suggesting recovery from AKI [37, 38] (Figure 2).

**Figure 2:**
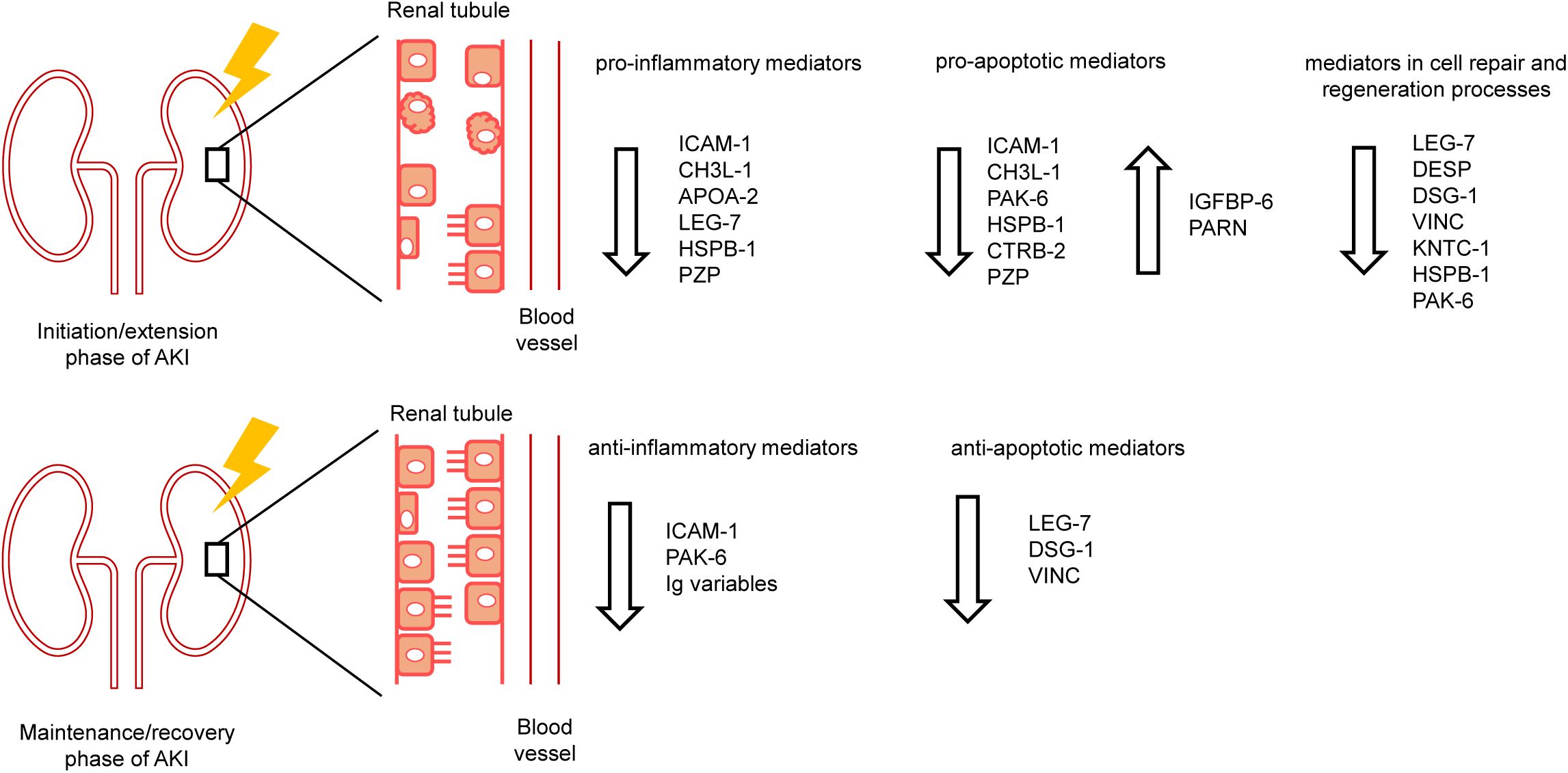
The role of candidate markers associated with immune responses, programmed cell death as well as processes of cell survival, repair and regeneration in the progression and recovery of AKI.

### Programmed cell death

RTECs are highly susceptible to apoptosis, which is characteristic of nephrotoxin-induced AKI and contributes to kidney dysfunction [12, 39, 40]. ICAM-1 participated in enriched pathways that negatively regulate extrinsic apoptotic signalling and endothelial cell apoptotic processes (Table 2). Additionally, CH3L-1 was reported to limit renal tubule and inflammatory cell apoptosis [41, 42, 43]. The decreased abundance of ICAM-1 and CH3L-1 in the AKI group suggested persistent activation of apoptotic pathways. HSPB-1 participates in enriched signalling pathways that negatively regulate oxidative stress-induced intrinsic apoptosis (Table 2) and PAK-6 contributes to anti-apoptotic signalling pathways [44] and their decreased abundance in the AKI group, suggested a lack of apoptotic inhibition and progression of AKI. IGFBP-6 participates in pro-apoptotic functions and its increased abundance in the AKI group contributes to the progression of AKI. Additionally, LEG-7 contributes to pro-apoptotic signalling pathways [45], DSG-1 participates in enriched pathways of apoptosis and apoptotic cleavage of cell adhesion proteins (Table 2), and VINC is regulator of apoptosis, and their decreased abundance in the AKI group suggested recovery of AKI. PARN inhibits p53-mediated apoptotic processes and had increased abundance in the AKI group, indicating recovery of AKI (Table 2). CTRB-2 and PZP participate in enriched serine-type endopeptidase activity and endopeptidase inhibition processes, respectively, and had decreased abundance in the AKI group, suggesting a reduction in their anti-inflammatory and anti-apoptotic activities [46] (Table 2) (Figure 2).

### Plasma lipoprotein assembly and cholesterol/lipid metabolism

Aberrant lipid metabolism is strongly associated with renal dysfunction and the progression of kidney diseases, including AKI. Lipid metabolism involves pathways that generate, store and transport lipids using different types of plasma lipoprotein particles in the blood [47, 48, 49]. The dysregulation of renal lipid metabolism is noted by declining HDL particles due to changes in lipid and apolipoprotein composition of HDL associated proteins, such as APOA-2, APOA-1, and lecithin-cholesterol acyltransferase (LCAT), disrupting reverse cholesterol transport, free fatty acid metabolism and HDL antioxidant functions associated with the kidneys [49, 50]. Additionally, ABC transporter proteins, such as ABCA-1 interacts with HDL apolipoproteins in reverse cholesterol transport. APOA-2 and ABCA-1 had decreased abundance in the AKI group, suggesting defective cholesterol metabolism in renal diseases and progression of AKI.

#### *Processes of cell survival*, *repair, and regeneration*

VINC, DSG-1, DESP, and LEG-7 participate in cell-cell and cell-ECM adhesion processes associated with the maintenance and recovery phases of AKI and had decreased abundance in the AKI group, suggesting disruption of repair and recovery of AKI. KNTC-1 prevents replicating cells from prematurely exiting mitosis [51], and its decreased abundance in the AKI group suggested disrupted tubule repair. HSPB-1 supports cell survival [52, 53] and PAK-6 participates in the MAPK signalling pathway [54] (Table 2), indicating that both candidate markers are involved in stress response and their decreased abundance in the AKI group, indicated inadequate cell survival and regeneration (Figure 2).

Machine learning (ML) models were explored to expand the search space for novel AKI candidate biomarkers in this study. The best performing ML model trained the XGBoost classifier and on average, 64% of patients with AKI were accurately assigned to the AKI group and 80% of patients without AKI to the non-AKI group. The ML model selected was better at predicting the control group than the disease group but had the best predictive value than the other ML models explored with the dataset provided.

Syntaxin-binding protein 5 (STXB-5), poly(A)-specific ribonuclease (PARN), and pregnancy zone protein (PZP) overlap the 20 ranked proteins used by the ML model to distinguish the study groups (Figure 1C), and the candidate markers identified using the Student’s t-test (Figure 1A). Fourteen of the remaining 17 proteins (Figure 1C) not identified as significantly differentially abundant by the Student’s t-test, profilin-1 (PROF-1) [58, 59], platelet factor 4 (PLF-4) [56, 57], beta-2-microglobulin (B2MG) [12], integrin alpha-IIb (ITA2B) [55], 14-3-3 protein zeta/delta (1433Z) [67, 68, 69], Ras-related protein Rap-1b-like protein (RP1BL) [70, 71], Transgelin-2 (TAGL2) [72, 73], Glyceraldehyde-3-phosphate dehydrogenase (G3P) [74], Zinc-alpha-2-glycoprotein (ZA2G) [75, 76], Tubulin beta chain (TBB5) [77], and Plexin-A2 (PLXA2) [78] have been associated with renal pathophysiological processes. This highlights the benefit of applying machine learning to proteomic data analysis in addition to traditional statistical testing.

Comparison of the urinary (u) and plasma (p) proteome profiles revealed that LEG-7, CH3L-1, IGFBP-6, HSPB-1, and KVD-33 overlapped both cohorts. Both pLEG-7 and uLEG-7 had decreased abundance in the AKI group compared to the non-AKI group and the fold-change of pLEG-7 was lower than that of uLEG-7. LEG-7 has been identified within RTECs and its upregulation is associated with maintenance phase of AKI progression, suggesting inadequate cell repair [60, 61, 62]. Additionally, pCH3L-1 had decreased abundance in the AKI group, while uCH3L-1 had increased abundance in the AKI group compared to the non-AKI group. The fold-change of pCH3L-1 was lower than that of uCH3L-1. The decreased abundance of pCH3L-1 in the AKI group is associated with recovery from AKI, whereas increased abundance of uCH3L-1 in the AKI group may be indicative of renal injury [42, 63, 64] (Figure 3).

**Figure 3:**
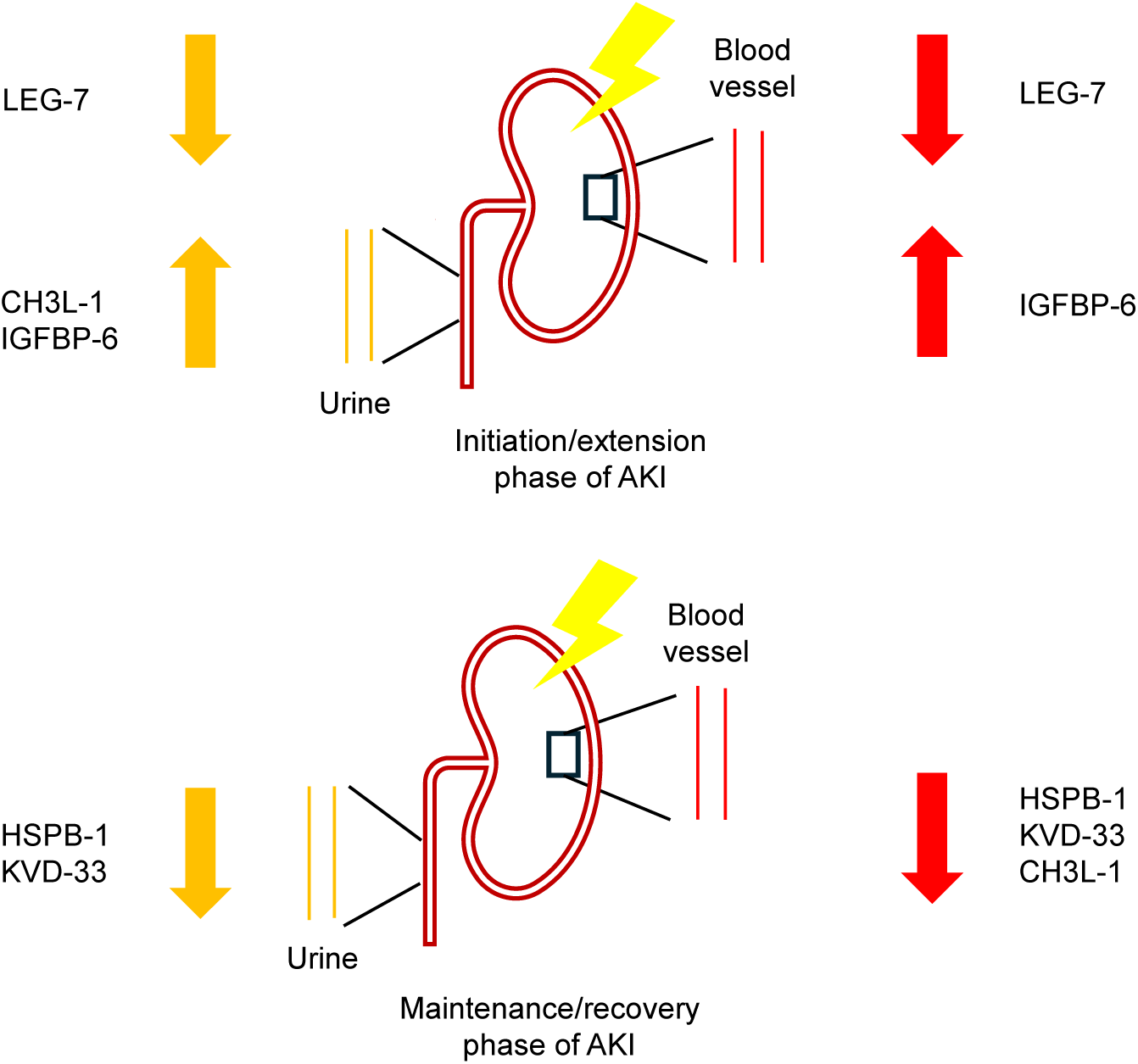
Comparison of overlapping candidate markers in urine (yellow) and plasma (red) in the progression and recovery of AKI.

Furthermore, pHSPB-1, pKVD-33, and pIGFBP-6 showed a similar fold-change to uHSPB-1, uKVD-33, and uIGFBP-6, respectively. Both pHSPB-1 and uHSPB-1, as well as pKVD-33 and uKVD-33 had decreased abundance in the AKI group, whereas pIGFBP-6 and uIGFBP-6 had increased abundance in the AKI group compared to the non-AKI group. The elevation of pHSPB-1 and uHSPB-1, as well as pKVD-33 and uKVD-33 occurs in response to renal tissue damage and their decline suggests renal recovery [52, 53, 79]. Lastly, increased abundance of pIGFBP-6 and uIGFBP-6 in the AKI group is associated with progression of renal dysfunction [80]. Comparison of candidate markers with dysregulated abundance levels provided evidence for the renal physiological dynamics in renal injury and disease (Figure 3).

## 5. Study limitations

The study was limited by the cohort design, as a longitudinal study design would be preferred for monitoring of AKI progression from a non-AKI state to an AKI state, allowing for early marker identification. Additionally, the large dynamic range of crude plasma protein abundances limits the identification of low abundant plasma proteins that can serve as putative protein biomarkers. Alternative sample preparation strategies may be employed to reduce the impact of high abundant plasma proteins on the depth of analysis.

## 6. Conclusion

This study enabled, for the first time, comprehensive identification of plasma proteome changes between TDF-exposed PLHIV, with and without AKI in the South African population. The identified candidate markers participated in the biological processes and pathways associated with progression and recovery of AKI, indicating their potential value as AKI biomarkers in PLHIV in South Africa. This work emphasizes the utility of machine learning in addition to conventional statistical methods in analysing proteomic data for the discovery of novel AKI biomarkers. The comparison of urinary and plasma proteome alterations provided insight into the physiological dynamics associated with these candidate markers in renal injury and disease, emphasizing the potential of combining various biomarkers from different biofluids in a panel for clinical monitoring of AKI and potentially other kidney diseases.

## Supplementary Materials

The following supplementary information is provided in this article: Supplementary Figure S1: Multi-scatter plots showing Pearson correlation of identified protein groups and peptides in the replicates of quality control samples. Supplementary Figure S2: Boxplot showing the median percentage coefficient of variation (% CV) for the replicates of quality control samples. Supplementary Table S1: Table of LC derived peak parameters. Supplementary Figure S3: Bar graph indicating the number of peptides and protein groups identified between the study groups. Supplementary Figure S4: Retrospective power analysis. Supplementary Table S2: Mean performance values for machine learning (ML) models evaluated. Supplementary Table S3: Mean performance values for machine learning (ML) models evaluated. Supplementary Table S4: Mean performance values for machine learning machine learning (ML) models evaluated. Supplementary Table S5: List of differentially abundant proteins generated from the AKI study.

## Data Availability Statement

The mass spectrometry proteomics data have been deposited to the ProteomeXchange Consortium via the PRIDE [82] partner repository with the dataset identifier PXD054218.

## Acknowledgements

The authors would like to acknowledge the individuals enrolled in this kidney injury study and thank the Perinatal HIV Research Unit (PHRU) for patient recruitment, sample collection and patient data management, as well as ReSyn Biosciences for the provision of the MagReSyn HILIC microspheres.

## Author Contributions

**R.J.M.** Conceptualization, data curation, formal analysis, investigation, methodology, project administration, validation, visualization, roles/writing - original draft. **S.F.** Supervision and writing - review and editing. **F.N.** Resources. **E.V.** Conceptualization and resources. **N.M.** Conceptualization, resources and funding acquisition. **I.S.G.** and **P.N.** Conceptualization, funding acquisition, methodology, project administration, supervision, validation and writing - review and editing.

## Corresponding authors address

Council for Scientific and Industrial Research (CSIR), NextGen Health Cluster, Building 20, Meiring Naude Road, Brummeria, Pretoria, 0001, South Africa.

