## Supplementary for "Plasma proteome profiling for AKI biomarker candidates associated with first-line ART in people living with HIV in South Africa"

### Supplementary Data

*Assessment of quality of LC-MS/MS data acquired*

The variability in the LC-MS/MS performance was evaluated over the course of five days by analysing Hela and Peptide_Pool QC replicates at the beginning of each day of analysis. Pearson correlations were performed, and a strong positive correlation was observed between the protein groups (≥ 0.93) and the peptides (≥ 0.875) identified in the Hela and Peptide_Pool QC replicates (Figure S1). A median percentage coefficient of variation (%CV) of 20,5% and 26,3% was observed, indicating the inter-day variation of the protein groups identified between the Hela and Peptide_Pool_QC replicates, respectively (Figure S2). To further assess the quality and reliability of the experimental data obtained, LC-derived peak parameters were evaluated. The peak capacity for all Hela and Peptide_Pool QC replicates was within the expected range for adequate peptide separation. As the peak capacity declines, the median full width half maximum (FWHM) of the peaks resolved in each replicate widens (indicated by an increased value in the median FWHM), reducing peak resolution. Additionally, the average data points per peak for the Hela and Peptide_Pool QC replicates were 6.1 and 7.1, respectively (Table S1).

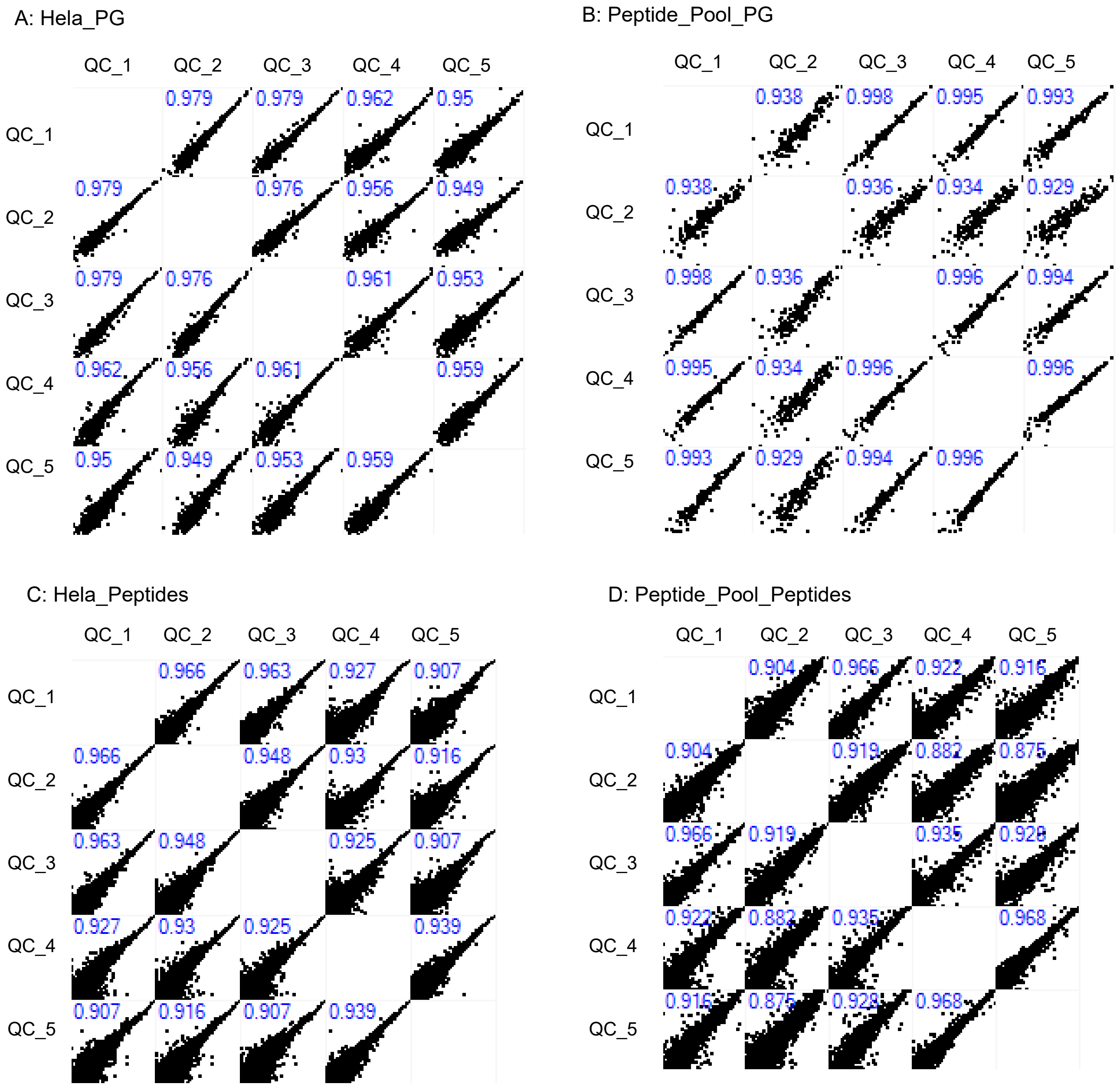

**Figure S1: Multi-scatter plots showing Pearson correlation of identified protein groups and peptides in Hela_QC (A, C) and Peptide_Pool_QC (B, D) replicates analysed over 5 days.**

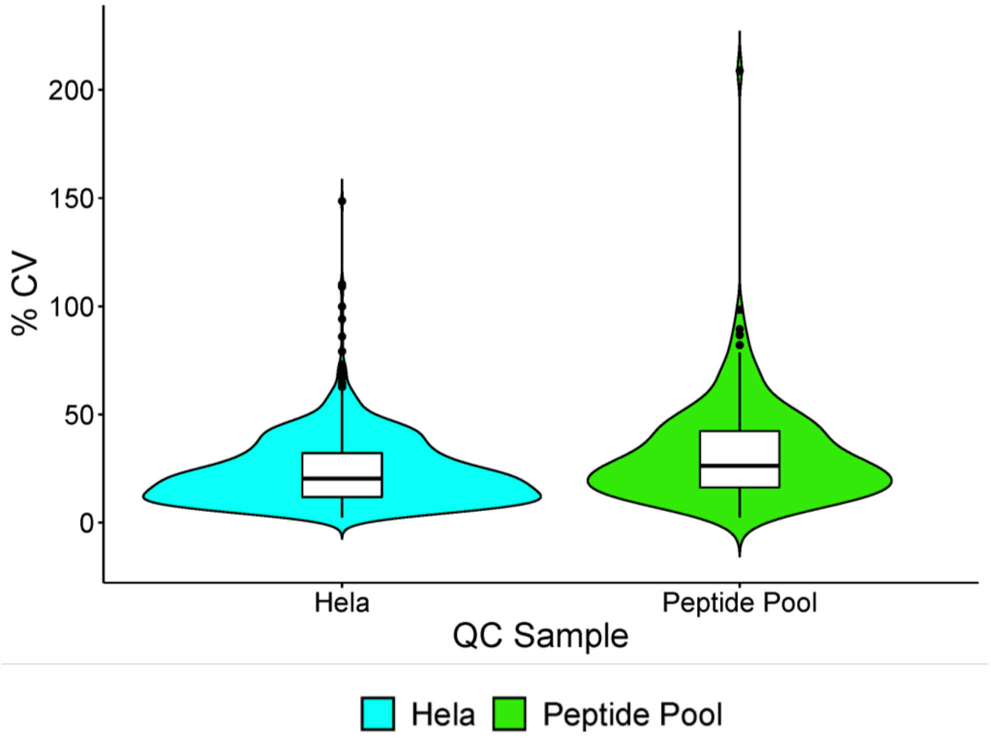

**Figure S2: Boxplot showing the median percentage coefficient of variation (% CV) for the Hela (A) in blue and Peptide_Pool (B) in green QC replicates.**

**Table S1: Table of LC derived peak parameters**. The average data points per peak, peak capacity and the median FWHM in minutes is tabulated for each Hela_QC and Peptide_Pool_QC replicate analysed.

| | Average data points per peak | Peak Capacity | Median FWHM (minutes) |
|---|---|---|---|
| **Hela_QC_1** | 5,9 | 154,85 | 0,0799 |
| **Hela_QC_2** | 5,98 | 152,53 | 0,0811 |
| **Hela_QC_3** | 5,92 | 146,12 | 0,0846 |
| **Hela_QC_4** | 6,54 | 107,51 | 0,1151 |
| **Hela_QC_5** | 6,18 | 98,33 | 0,1258 |
| **Average data points for Hela** | 6,10 | | |
| **Peptide_Pool_QC_1** | 6,57 | 178,49 | 0,0693 |
| **Peptide_Pool_QC_2** | 6,88 | 174,35 | 0,0709 |
| **Peptide_Pool_QC_3** | 6,62 | 170,98 | 0,0723 |
| **Peptide_Pool_QC_4** | 7,88 | 125,33 | 0,0987 |
| **Peptide_Pool_QC_5** | 7,52 | 118,98 | 0,104 |
| **Average data points for Peptide_Pool** | 7,09 | | |

*Overview of AKI cohort*

Within the individual study conditions, an average of 3 365 modified peptides and 380 protein groups were identified in the AKI group, while an average of 3 113 modified peptides and 352 protein groups were identified in the non-AKI group.

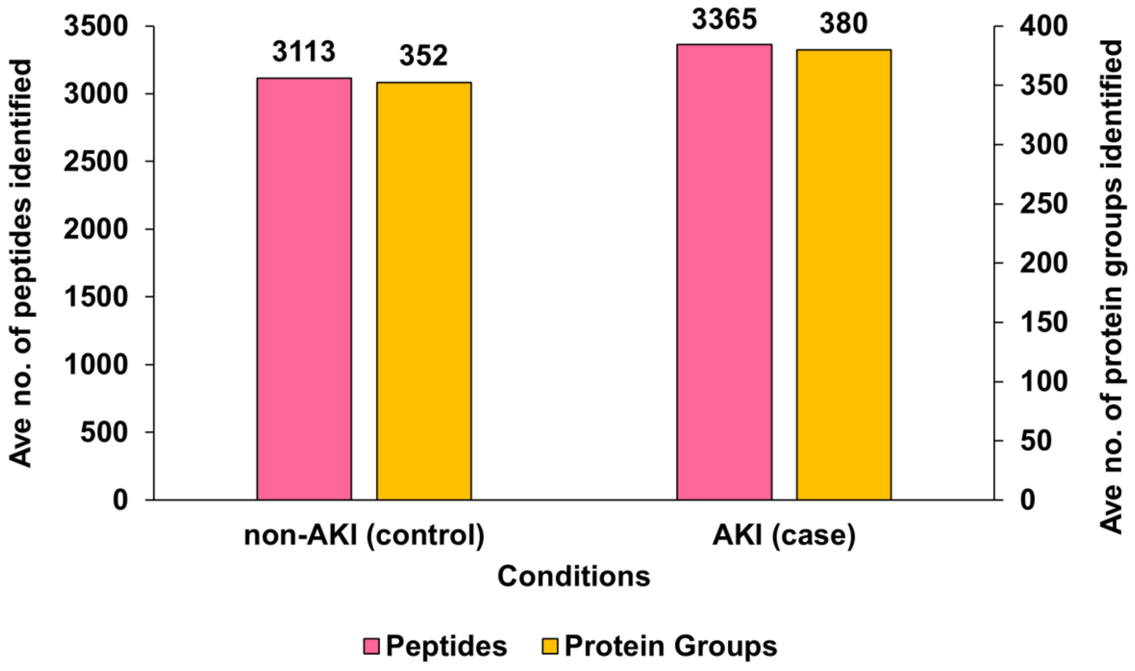

**Figure S3: Bar graph indicating the number of peptides (pink) and protein groups (orange) identified between the AKI and non-AKI groups.**

*Retrospective power analysis*

The retrospective power analysis performed showed that at a power of 80%, the appropriate fold-change cut-off for the plasma candidate markers identified in the AKI study is ~ 1,68. The significance criterion for the differentially abundant proteins identified was q-value ≤ 0.05, $\log_2 FC \geq 0.75$ ($FC \geq 1.68$).

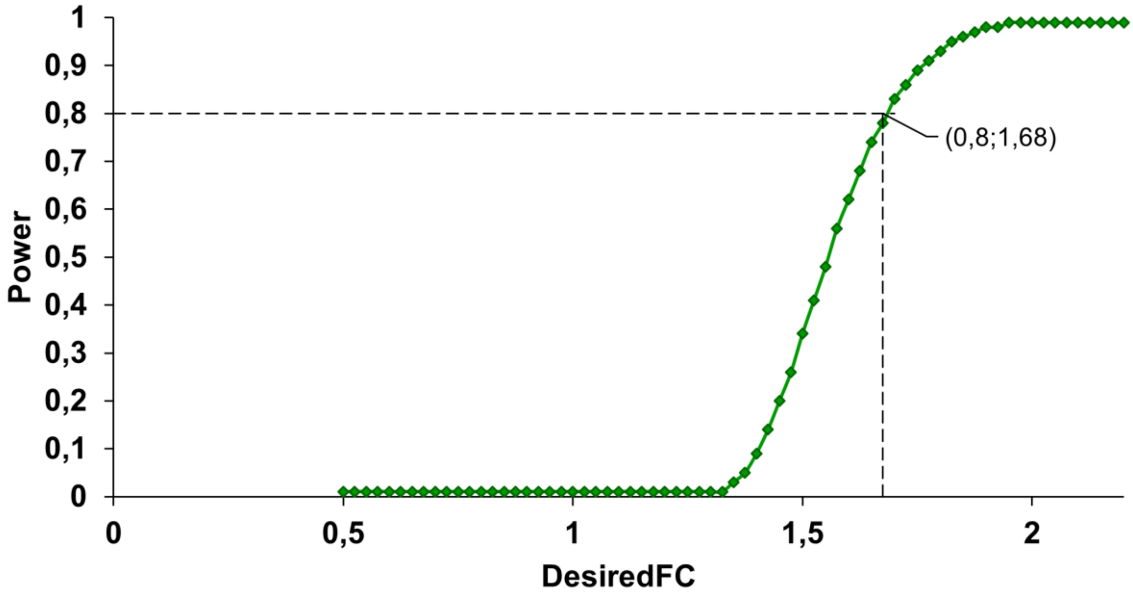

**Figure S4: Line graph indicating the estimated power of the AKI study and the corresponding desired fold-change (FC) defining the candidate markers**. The desired FC is given at an FDR of 1%. Power analysis completed using MSstats (v3.4.1) in RStudio.

*Exploring the performance of machine learning models*

The performance of the different machine learning (ML) models explored was evaluated based on the scores (0 – 1; 0: poor performance and 1: excellent performance) given for accuracy, precision, recall, F1_score, ROC_AUC and PR_AUC.

The performance of the 5 feature selection methods was evaluated using the XGBoost classifier with a 'no imputation' method. The 'ExtraTrees' option was preferred to the 'None' option, as the 'None' option can create bias in the important protein features identified by the selected ML model(s). The performance scores for each ML model were reported in Table S1.

Secondly, the performance of 7 ML classifiers was assessed using the 'ExtraTrees' feature selection method and the 'zero imputation' method, where 'zero' was imputed in the place of missing values in the dataset. The XGBoost classifier had the best performance across all classifiers evaluated. The performance scores for each ML model were reported in Table S2.

Lastly, the performance of the 'no', 'zero', as well as the 'median' imputation methods was evaluated using the 'ExtraTrees' feature selection method and the XGBoost classifier. The 'no imputation' method was preferred to the 'median imputation' method, to ensure that no bias or loss of information from the features with missing values occurs. The performance scores for each ML model were reported in Table S3.

**Table S2: Mean performance values for ML models evaluated in Figure S5A.** ML models were built using the ‘no’ imputation strategy and XGBoost classifier to evaluating various feature selection methods (ExtraTrees, k-best [mutual_info_classif, chi2 and f-classif] and none) in the ML pipelines for the AKI study.

| | ExtraTrees | k-best (mutual_info_classif) | k-best (f_classif) | k-best (chi2) | None | Best Score | Best Score (excl. 'none') |
|---|---|---|---|---|---|---|---|
| **Accuracy** | 0,7363 | 0,7304 | 0,7215 | 0,7126 | 0,7578 | **None** | **ExtraTrees** |
| **Precision** | 0,6884 | 0,6839 | 0,6601 | 0,6428 | 0,7223 | **None** | **ExtraTrees** |
| **Recall** | 0,6360 | 0,6169 | 0,6185 | 0,6342 | 0,6385 | **None** | **ExtraTrees** |
| **F1 Score** | 0,6485 | 0,6375 | 0,6276 | 0,6269 | 0,6671 | **None** | **ExtraTrees** |
| **ROC_AUC** | 0,8154 | 0,7985 | 0,8082 | 0,7604 | 0,8327 | **None** | **ExtraTrees** |
| **PR_AUC** | 0,7360 | 0,7175 | 0,7181 | 0,6515 | 0,7515 | **None** | **ExtraTrees** |

**Table S3: Mean performance values for ML models evaluated in Figure S5B.** ML models were built using the ‘zero’ imputation strategy and the ExtraTrees feature selection method to explore the various ML classifiers (AdaBoost, LogisticRegression, RandomForest, k-NeighboursClassifier, DecisionTree and LinearSVC) in the ML pipelines for the AKI study.

| | AdaBoost | LogisticRegression | RandomForest | k-Neighbours Classifier | DecisionTree | LinearSVC | XGBoost | Best Score |
|---|---|---|---|---|---|---|---|---|
| **Accuracy** | 0,7207 | 0,7119 | 0,7215 | 0,6074 | 0,6578 | 0,7074 | 0,7244 | **XGBoost** |
| **Precision** | 0,6667 | 0,6455 | 0,6766 | 0,0000 | 0,5774 | 0,6452 | 0,6732 | **RandomForest** |
| **Recall** | 0,6153 | 0,6162 | 0,5867 | 0,0000 | 0,5425 | 0,5338 | 0,5924 | **LogisticRegression** |
| **F1 Score** | 0,6263 | 0,6193 | 0,6155 | 0,0000 | 0,5505 | 0,5668 | 0,6211 | **AdaBoost** |
| **ROC_AUC** | 0,7859 | 0,7942 | 0,8087 | 0,7076 | 0,6371 | 0,7994 | 0,8102 | **XGBoost** |
| **PR_AUC** | 0,6747 | 0,6684 | 0,7105 | 0,6836 | 0,6496 | 0,7064 | 0,7183 | **XGBoost** |

**Table S4: Mean performance values for ML models evaluated in Figure S5C.** ML models were built using the ExtraTrees feature selection method with the XGBoost classifier to evaluate the 'no', 'zero' and 'median' imputation strategies in the ML pipelines for the AKI study.

| | No imputation | Zero imputation | Median imputation | Best Score |
|---|---|---|---|---|
| **Accuracy** | 0,7363 | 0,7244 | 0,7370 | **Median imputation** |
| **Precision** | 0,6884 | 0,6732 | 0,6883 | **No imputation** |
| **Recall** | 0,6360 | 0,5924 | 0,6329 | **No imputation** |
| **F1 Score** | 0,6485 | 0,6211 | 0,6507 | **Median imputation** |
| **ROC_AUC** | 0,8154 | 0,8102 | 0,8239 | **Median imputation** |
| **PR_AUC** | 0,7360 | 0,7183 | 0,7504 | **Median imputation** |

**Table S5: List of differentially abundant proteins generated from the AKI study.** Spectronaut (v17) generated 34 candidate markers in the AKI study and are arranged in descending order of average log2 ratio (fold-change) at q-value < 0.05.

| Protein Names | Protein Descriptions | Average Log2 Ratio (fold change) |
|---|---|---|
| **RIMS-3** | Regulating synaptic membrane exocytosis protein 3 | 1,10 |
| **PARN** | Poly(A)-specific ribonuclease PARN | 0,97 |
| **IGFBP-6** | Insulin-like growth factor-binding protein 6 | 0,90 |
| **KV-116** | Immunoglobulin kappa variable 1-16 | -0,78 |
| **AIG-1** | Androgen-induced gene 1 protein | -0,78 |
| **KVD-33** | Immunoglobulin kappa variable 1D-33 | -0,81 |
| **STXB-5** | Syntaxin-binding protein 5 | -0,84 |
| **DYH-9** | Dynein axonemal heavy chain 9 | -0,85 |
| **LV-208** | Immunoglobulin lambda variable 2-8 | -0,87 |
| **SIG-16** | Sialic acid-binding Ig-like lectin 16 | -0,88 |
| **CH3L-1** | Chitinase-3-like protein 1 | -0,89 |
| **AL3A-2** | Aldehyde dehydrogenase family 3 member A2 | -0,89 |
| **PZP** | Pregnancy zone protein | -0,94 |
| **HV-364** | Immunoglobulin heavy variable 3-64 | -0,98 |
| **TFP-11** | Tuftelin-interacting protein 11 | -1,06 |
| **LV-211** | Immunoglobulin lambda variable 2-11 | -1,09 |
| **ICAM-1** | Intercellular adhesion molecule 1 | -1,10 |
| **PAK-6** | Serine/threonine-protein kinase PAK 6 | -1,12 |
| **KNTC-1** | Kinetochore-associated protein 1 | -1,14 |
| **WRP-73** | WD repeat-containing protein WRAP73 | -1,15 |
| **HSPB-1** | Heat shock protein beta-1 | -1,23 |
| **HV-374** | Immunoglobulin heavy variable 3-74 | -1,33 |
| **LV-552** | Immunoglobulin lambda variable 5-52 | -1,42 |
| **DESP** | Desmoplakin | -1,43 |
| **KVD-37** | Probable non-functional immunoglobulinn kappa variable 1D-37 | -1,46 |
| **DSG-1** | Desmoglein-1 | -1,47 |
| **CTRB-2** | Chymotrypsinogen B2 | -1,55 |
| **APOA-2** | Apolipoprotein A-II | -1,70 |
| **VINC** | Vinculin | -2,15 |
| **ABCA-1** | Phospholipid-transporting ATPase ABCA1 | -2,18 |
| **IGHG-1** | Immunoglobulin heavy constant gamma 1 | -2,52 |
| **LV-545** | Immunoglobulin lambda variable 5-45 | -2,81 |
| **LV-312** | Immunoglobulin lambda variable 3-12 | -3,00 |
| **LEG-7** | Galectin-7 | -3,27 |
